# *NRG1* is required for the function of the TIR-NLR immune receptors Roq1 and RPP1 in *Nicotiana. benthamiana*

**DOI:** 10.1101/284471

**Authors:** Tiancong Qi, Alex Schultink, Julie Pham, Myeong-Je Cho, Brian J. Staskawicz

## Abstract

The plant immune system involves a large family of nucleotide-binding leucine-rich repeat (NLR) intracellular immune receptors. These immune receptors often function to directly or indirectly mediate the perception of specific pathogen effector proteins secreted into the cell. Activation of these immune receptors typically results in activation of the immune system and subsequent suppression of pathogen proliferation. Although many examples of NLR receptors are known, a mechanistic understanding of how receptor activation ultimately leads to an immune response is not well understood. A subset of the NLR proteins contain a TIR domain at their N terminus (TNL). One such TNL, the N gene, was previously shown to depend on a non-TIR NLR protein, N requirement gene 1 (NRG1) for immune function. We tested additional NLR proteins in *Nicotiana benthamiana* for dependency on NRG1. We found that two additional TIR-NLR proteins, Roq1 and RPP1, also require NRG1 but that two coiled-coil NLR proteins, Bs2 and Rps2, do not. This finding suggests that NRG1 may be a conserved component of TNL signaling pathways.

## Introduction

Plants have immune systems to recognize potential virulent pathogens. In response to pathogen invasion, pattern recognition receptors of plants are able to recognize conserved pathogen-associated molecular patterns (PAMPs), such as flagellin, and initiate pattern triggered immunity (PTI) to protect plants from pathogen infection (Monaghan and Zipfel, 2012; Tang et al., 2017; Zipfel, 2014). Gram-negative bacterial pathogens employ type III secretion systems (TTSS) to directly inject effector proteins into host cells that result in the suppression of plant defense, causing disease (Xin and He, 2013).

Host plants employ intracellular nucleotide-binding domain and leucine-rich repeat (NLR) proteins to recognize pathogenic effectors and activate effector-triggered immunity (ETI) (Dangl, 2013; Jones et al., 2016; Khan et al., 2016; Chisholm et al., 2006). ETI can lead to programmed cell death, known as the hypersensitive response (HR), at the infection site which is thought to suppress pathogen proliferation (Cui et al., 2015).

The N-terminus of NLR proteins usually contains a coiled-coil (CC) domain or a Toll/interleukin-1 receptor (TIR) domain, which classify the NB-LRR proteins into (CNL) proteins and (TNL) proteins (Zhang et al., 2017; Qi and Innes, 2013). The CNL protein Bs2 which was identified from pepper recognizes the effector AvrBs2 from *Xanthomonas euvesicatoria* (Xcv) and confers resistance to Xcv in pepper and tomato (Kearney and Staskawicz, 1990; Tai et al., 1999). *Arabidopsis* Rps2 is a CNL protein that recognizes the effector AvrRpt2 from *Pseudomonas syringae* and activates ETI responses (Mindrinos et al., 1994; Bent et al., 1994). The *Arabidopsis* TNL receptor Recognition of *Peronospora parasitica* 1 (RPP1) activates resistance responses after binding to the effector Arabidopsis Thaliana Recognized 1 (ATR1) from *Hyaloperonospora arabidopsidis* and induces an HR response in both *Arabidopsis* and *N. benthamiana* (Rehmany, 2005; Steinbrenner et al., 2015; Schreiber et al., 2016). The tobacco TNL protein N binds to the helicase fragment (p50) of tobacco mosaic virus (TMV) and triggers HR response and resistance to TMV (Erickson et al., 1999). Virus induced gene silencing (VIGS) of *N requirement gene 1* (*NRG1*) compromised N-mediated resistance to TMV in both *Nicotiana benthamiana* and *Nicotiana edwardsonii*. NRG1 is a CNL rather than a TNL protein, and so far was only found to be required for N-mediated resistance to TMV (Peart et al., 2005). The molecular events that specify recognition and activation of various NLR proteins in the ETI signaling pathways remain largely unknown.

The bacterial pathogens from the genera *Xanthomonas* and *Pseudomonas* cause severe diseases in various plants. The ability of a particular species or pathovar of *Xanthomonas* or *Pseudomonas* to cause disease in a plant is often dependent on the specific repertoire of TTSS effector proteins present in the pathogen (Keen, 1990; Collmer et al., 1999). *N. benthamiana* is resistant to species of *Xanthomonas* and *Pseudomonas* that contain the homologous effector proteins XopQ and HopQ1 from *Xanthomonas* and *Pseudomonas* respectively (Schwartz et al., 2015; Wei et al., 2007) Perception of XopQ/HopQ1 in *N. benthamiana* is mediated by the TNL protein Roq1 (Schultink et al). Like other TNL proteins, Roq1-mediated resistance also depends on the presence of Enhanced Disease Susceptibility 1 (EDS1) (Schultink et al., 2017; Adlung and Bonas, 2017; Adlung et al., 2016).

The TNL protein Recognition of XopQ 1 (Roq1) was identified as being genetically required for perception of XopQ/HopQ1 in *N. benthamiana*, and was shown to interact with XopQ (Schultink et al., 2017). In this study, we identified NRG1 as being required for the function of Roq1. *N. benthamiana* mutants of NRG1 and Roq1 were generated using the CRISPR/Cas9 system to facilitate the study of this immune pathway. Immune activation and disease resistance in response to XopQ/HopQ1 are absent in the *nrg1* and *roq1* mutants, as previously observed for the *eds1*. The NRG1 was found to be required for the function of an additional TNL but not two CNL proteins, suggesting that NRG1 may be a conserved component of TNL immune signaling.

## Results

### Virus-induced gene silencing of *NRG1* and *EDS1* compromised Several TNLs –mediated HR response in *N. benthamiana*

To investigate the role of *NRG1* and *EDS1* in ETI pathway, we firstly adopted virus-induced gene silencing (VIGS) (Liu et al., 2002) strategy to silence *NRG1* and *EDS1* in *N. benthamiana,* and observed hypersensitive response (HR) induced by *Agrobacterium*-mediated transient expression of several TNLs and their recognized effectors, including XopQ, HopQ1, RPP1-ATR1 and N-P50, and CNLs and their cognate elicitors, including Bs2-AvrBs2.While activation of Roq1 by *Agrobacterium*-mediated expression of XopQ leads to a strong HR in *N. tabacum*, it often leads to only a mild chlorotic response in *N. benthamiana* under typical conditions (Schultink et al., 2017; Adlung and Bonas, 2017). We observed that keeping *N. benthamiana* leaf tissue in the dark after infiltration enhanced the effector-triggered HR response, so *N. benthamiana* leaves were covered in aluminum foil during the following experiments for observation of HR phenotype.

Previously results showed that *N*-mediated resistance to TMV was abolished in *EDS1* silencing plant in *N. benthamiana* (Liu et al., 2002), and XopQ/HopQ1 effectors-caused HR was abolished in *eds1* knockout mutant in *N. benthamiana* (Adlung et al., 2016). Consistently, our results also showed that silencing of *EDS1* disrupted HR phenotype caused by N-P50, XopQ, HopQ1, and also RPP1-ATR1 (Supplemental Figure 1). HR phenotype caused by Bs2 and AvrBs2, a CNL protein and its recognized effector, was not affected in *EDS1* silencing plant (Supplemental Figure 1), consistent with previously well understand knowledge that *EDS1* is required for multiple TNLs-mediated, but not for CNLs-mediate ETI response.

**Figure 1.**
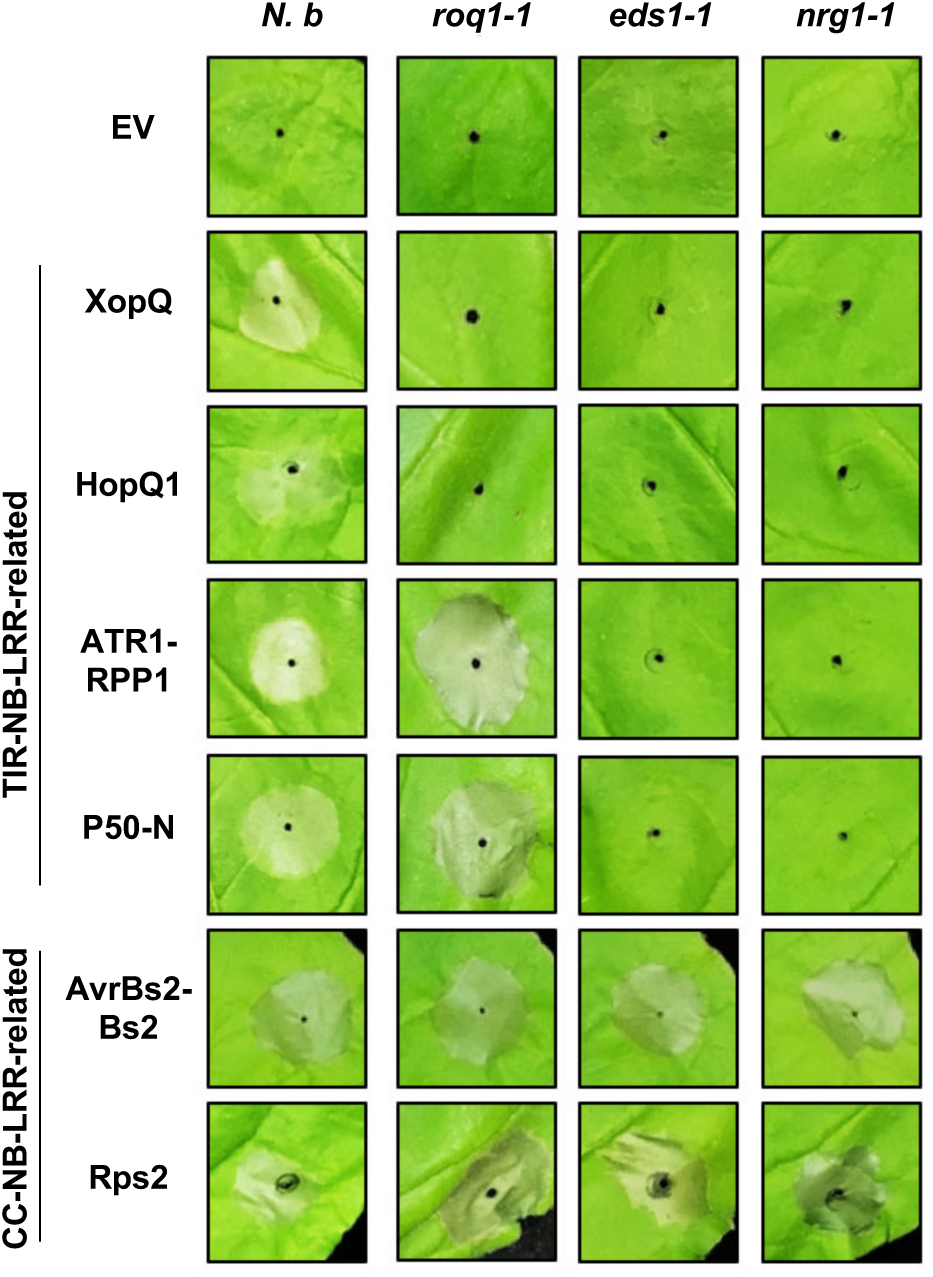
Roles of Roq1, EDS1 and NRG1 in effector-triggered hypersensitive response. Phenotypes of hypersensitive response (HR) in leaves of N. benthamiana wild-type (N. b), roq1-1, eds1-1, or nrg1-1 with Agrobacterium-mediated transient expression of the empty vector control (EV), XopQ, HopQ1, ATR1-RPP1, N-P50, AvrBs2-Bs2, or Rps2. The infiltrated leaves were wrapped up with aluminum foil and images were taken 2 days post-infiltration

NRG1 was previously demonstrated to be required for N-mediated resistance to TMV in *N. benthamiana* (Peart et al., 2005). N is a TNL protein that can recognize p50 from TMV and activate ETI pathway (Erickson et al., 1999). Consistently, the plants silenced for *NRG1* also showed compromised N-activated HR response (Supplemental Figure 1). More interestingly, we found that *NRG1* silenced plants also failed to have an HR phenotype in response to the expression of XopQ, HopQ1, or RPP1-ATR1 (Supplemental Figure 1). The Bs2-AvrBs2-activated HR was not affected in the plants silenced for *NRG1* (Supplemental Figure 1). The results imply that NRG1 may be a conserved component in TNLs-mediated ETI signaling pathway.

### Generation of *roq1* and *nrg1* mutants in *N. benthamiana* using CRISPR/Cas9

Stable knockout mutants are important for investigating plant immune responses. In order to have stable mutants of *Roq1, NRG1* and *EDS1* for investigating their roles in XopQ/HopQ1-triggered immunity, we obtained *eds1* mutants from our previous study (Schultink et al., 2017), and constructed CRISPR/Cas9-mediated mutations into *Roq1* (Supplemental Figure 2A-C) and *NRG1* (Supplemental Figure 2D-G) in *N. benthamiana* (Jinek et al. 2012; Schultink et al. 2017). Two functional guide RNAs targeting *Roq1* and two functional guide RNAs targeting *NRG1* were used to generate stable transformants in *N. benthamiana*. Independent homozygous lines, including frameshift alleles of deletion and insertion, were obtained through selfing and named *roq1-1, roq1-2, nrg1-1, nrg1-2*, and *nrg1-3* (Supplemental Figure 2).

**Figure 2.**
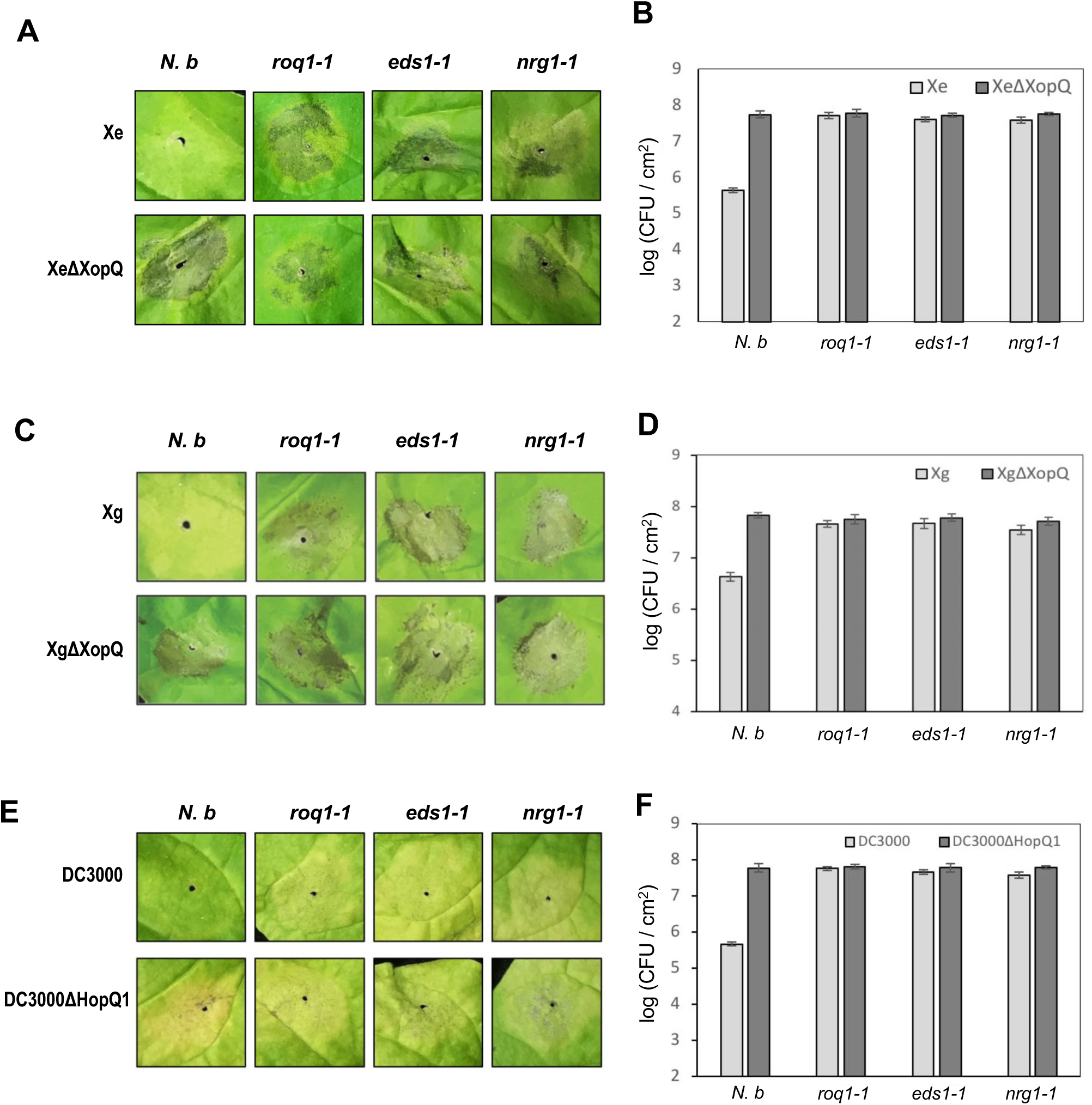
Roq1, EDS1 and NRG1 are required for plant resistance to bacterial pathogens. (A and B) Disease symptoms (A) and bacterial populations (B) in leaves of N. benthamiana wild-type, roq1-1, eds1-1, or nrg1-1 after syringe infiltration with Xanthomonas euvesicatoria (Xe) and the XopQ knockout (XeΔXopQ) at low inoculum (OD600 = 0.0001). Disease symptoms were recorded at 12 days post-infiltration, and bacterial growth was assayed 6 days post-infiltration. (C and D) Disease symptoms (C) and bacterial populations (D) in leaves of N. benthamiana wild-type, roq1-1, eds1-1, or nrg1-1 after syringe infiltration with Xanthomonas gardneri (Xg). and the XopQ knockout (XgΔXopQ) at low inoculum (OD600 = 0.0001). Disease symptoms were recorded at 10 days post-infiltration, and bacterial growth was assayed 6 days post-infiltration. (E and F) Disease symptoms (E) and bacterial populations (F) in leaves of N. benthamiana wild-type, roq1-1, eds1-1, or nrg1-1 after syringe infiltration with Pseudomonas syringae pv. tomato DC3000 and the HopQ1 knockout (DC3000ΔHopQ1) at low inoculum (OD600 = 0.0001). Disease symptoms were recorded at 8 days post-infiltration. Bacterial growth was assayed 5 days post-infiltration.

### *Roq1, NRG1* and *EDS1* play key roles in hypersensitive response of ETI pathway

To better investigate the roles of *Roq1, NRG1* and *EDS1* in ETI pathway, we further observed HR response in the *N. benthamiana* wild-type, *roq1, nrg1* and *eds1* mutants after *Agrobacterium*-mediated transient expression of several TNLs, CNLs and/or their recognized effectors.

As shown in Figure 1, *Agrobacterium*-mediated transient expression of XopQ, HopQ1, RPP1-ATR1, N-P50, Bs2-AvrBs2 and Rps2 in *N. benthamiana* wild-type resulted in an HR. Consistent with our previous study that Roq1 mediates the perception of XopQ (Schultink et al., 2017), XopQ and its close homolog HopQ1 were unable to trigger HR in *roq1* mutants (Figure 1 and Supplemental figure 3). Similar with wild-type, *roq1* mutants exhibited HR in response to the expression of Bs2-AvrBs2, Rps2, N-P50, and RPP1-ATR1 (Figure 1 and supplemental Figure 3), demonstrating that Roq1 is not required for these other immune pathways.

Consistent with our *NRG1* and *EDS1* silencing plant phenotypes, all the TNLs and/or cognate elicitors we tested, including XopQ, HopQ1, N-p50 and RPP1-ATR1, were unable to trigger HR in the *nrg1* and *eds1* mutants, while the CNLs and recognized effectors (Bs2-AvrBs2 and Rps2) could (Figure 1 and supplemental Figure 3), further implying that both EDS1 and NRG1 are particularly conserved components for TNL-activated HR, but not CNL-regulated HR.

Taken together (Figure 1, supplemental Figure 1 and 3), these data suggested that Roq1 specially mediates XopQ/HopQ1-triggered HR, and that EDS1 and NRG1 are most likely conserved components in TNLs-mediated ETI pathway.

### *Roq1, NRG1* and *EDS1* function in effector-triggered resistance to TMV

A TMV-based viral replicon system containing green fluorescent protein (GFP) (Marillonnet et al., 2005), has previously been used as a reporter for plant immune activation (Schultink et al 2017). With this system, GFP is strongly expressed from the viral replicon in the absence of ETI but expression of GFP is suppressed if ETI is activated by co-expression of an immune elicitor. The transiently expressed TMV replicon containing GFP (TMV-GFP) proliferated well in the leaves of *N. benthamiana* wild-type harboring no N gene (Supplemental Figure 4). The transient expression of Bs2-AvrBs2, RPP1-ATR1 or XopQ1 triggered HR in *N. benthamiana* wild-type and reduced expression of GFP (Supplemental Figure 4), indicating that effector-triggered immunity is able to confer resistance to TMV in *N. benthamiana*.

XopQ expression was unable to trigger HR in the *roq1* mutant plants, and TMV-GFP proliferate well, whereas the expression of Bs2-AvrBs2 or RPP1-ATR1 induced HR and blocked TMV-GFP proliferation (Supplemental Figure 4). The expression of TNL-related RPP1-ATR1 or XopQ could no longer trigger HR in both *eds1* and *nrg1* mutants, and TMV-GFP was able to proliferate in *eds1* and *nrg1* mutants. On the other hand, the CNL-related Bs2-AvrBs2 could trigger HR and prevented TMV-GFP expression in the *eds1* and *nrg1* mutants (Supplemental Figure 4).

These results (Supplemental Figure 4) indicated that Roq1 activates ETI and confers defense against TMV specifically in response to XopQ, whereas NRG1 and EDS1 probably contribute to TNL-activated resistance to TMV, but not CNL-activated immunity.

### NRG1 is essential for suppressing the growth of *Xanthomonas* and *Pseudomonas* in *N. benthamiana*

Having shown that NRG1 is required for Roq1-mediated HR, we carried out bacterial growth assays to examine whether this protein is also required for Roq1-mediated disease resistance to bacterial pathogens from the genera *Xanthomonas* and *Pseudomonas*.

Consistent with previous results, *Xanthomonas euvesicatoria* (Xe) grew much higher and displayed stronger disease phenotypes in *eds1* mutant compared with *N. benthamiana* wild type, and Xe grew dramatically less than the XopQ knockout (XeΔXopQ) (Figure 2A-B) (Schultink et al., 2017; Adlung et al., 2016). Furthermore, we found that the growth of Xe and XeΔXopQ on *roq1* and *nrg1* was similar with XeΔXopQ on wild-type plants, and all the mutants displayed disease symptoms (Figure 2A-B), demonstrating that *roq1* and *nrg1* mutants cannot respond to XopQ, and fail to activate defense response (Figure 2A-B). Similar results were observed when we inoculated wild-type plants, *roq1, eds1* and *nrg1* with *Xanthomonas gardneri* (Xg), the XopQ knockout (XgΔXopQ), *Pseudomonas syringae* pv. *tomato* DC3000 and the HopQ1 knockout (DC3000ΔHopQ1) (Figure 2C-F). The Xg and DC3000 grew dramatically more on *roq1, nrg1* and *eds1* than on the *N. benthamiana* wild-type, and the knockouts (XgΔXopQ and DC3000ΔHopQ1) grew similar amount in wild-type, *roq1, nrg1* and *eds1* (Figure 2C-F).

Taken together (Figure 2), these results demonstrated that the *roq1, nrg1* and *eds1* mutants fail to activate XopQ/HopQ1-triggered ETI and were more susceptible to *Xanthomonas* and *Pseudomonas*, and that Roq1, EDS1 and NRG1 are collectively required and probably function in the same ETI pathway triggered by XopQ/HopQ1.

## Discussion

Plants employ different classes of NLR immune receptors that recognize pathogen effectors to trigger ETI and protect themselves from pathogen invasion (Gouveia *et al.*, 2017; Cui *et al.*, 2015; Rajamuthiah and Mylonakis, 2014). Elucidation of the molecular mechanisms controlling ETI will be necessary to develop novel strategies for disease control and crop breeding. The bacterial pathogens *Xanthomonas* and *Pseudomonas* take advantage of their effectors including XopQ and HopQ1, to infect host plants and cause disease (Ferrante et al 2009). The TNL protein Roq1 was previously identified as an immune receptor meditating recognition of XopQ/HopQ1 in *N. benthamiana* (Schultink et al., 2017). In this study, we found that the CNL protein NRG1 is also required for perception of XopQ/HopQ1. Stable mutants of *roq1, nrg1* and *eds1* allowed for detailed investigation into the roles of these three genes in plant immunity. These genes are essential for XopQ/HopQ1-triggered HR and disease resistance. As previously shown for EDS1, we observed that NRG1 is required for additional TNL-activated HR and defense, such as N, RPP1 and Roq1, but not for CNL-mediated HR and defense. It seems that EDS1 and NRG1 act as central component and integrate different signals from multiple effectors/TNLs to a single ETI pathway.

## Materials and Methods

### Generation of *N. benthamiana* mutants and plant growth conditions

Guide sequences were designed to target the *Roq1* and *NRG1* genes and corresponding oligonucleotides were annealed and ligated into a pDONR-based entry plasmid containing the *CAS9* gene and a U6-26 promoter for guide expression (Schultink et al., 2017). An LR reaction (Life Technologies) was used to move the guide and CAS9 cassette into a Gateway-compatible version of the pCambia2300 vector. These constructs were transformed into the *Agrobacterium* strain GV3101. Stable *N. benthamiana* transformants were generated by *Agrobacterium*-mediated transformation. The primers used for vector construction and genotyping are listed in Supplemental Table 1. The *eds1-1* mutant was previously described (Schultink et al., 2017). The *N. benthamiana* wild-type and mutants were grown under a 16-h (25 to 28°C)/8-h (22 to 25°C) light/dark photoperiod condition.

### Transient expression using *Agrobacterium tumefaciens*

The coding sequences of XopQ, HopQ1, Bs2, AvrBs2, N, P50, ATR1, RPP1 were fused with C-terminal 3flag and constructed into PE1776 vector. The Rps2 with HA tag were described previously (Day, 2005). The TMV reporter with fusion of green fluorescent protein (TMV-GFP) were described previously (Marillonnet et al., 2005). *Agrobacterium tumefaciens* containing the vectors were shaken, harvested, resuspended in infiltration buffer (10 mM MES, 10 mM MgCl_2_, and 0.2 mM acetosyringone), and co-infiltrated into leaves of the *N. benthamiana* wild-type, the *roq1, eds1*, and *nrg1* mutants using a needleless syringe. For figure 1 and Supplemental Figure 1, 3 and 4, the infiltrated leaves were wrapped up with aluminum foil for growth under dark. The leaves were observed and imaged 2-3 days post infiltration under white light or long-wave ultraviolet. Three biological repeats were performed for each experiment.

### *In planta* bacterial-growth assays

The bacteria *Pseudomonas syringae pv.* tomato DC3000, the HopQ1 knockout (DC3000ΔhopQ1), *Xanthomonas euvesicatoria* (Xe), *Xanthomonas gardneri* (Xg) and the XopQ knockouts (XeΔXopQ and XeΔXopQ) were grown at 28°C overnight in liquid NYG (0.5% peptone, 0.3% yeast extract, 2% glycerol) with 100 µg/mL rifampicin, harvested, resuspended in 10 mM MgCl_2_ buffer (OD_600_=0.0001), and infiltrated into the leaves of *N. benthamiana* wild-type and mutants. Leaf punches were collected at 5 or 6 days post-infiltration, homogenized, diluted and plated on NYG solid medium with 100 µg/mL rifampicin. Colonies were counted 2-3 days after plating. Three biological repeats were performed for each experiment.

### Virus induced gene silencing

A 318bp fragment of *NRG1* was inserted into the TRV2 VIGS vector (Liu et al., 2002). The primer sequences are listed in Supplemental Table S1. The TRV2 vectors for silencing *GUS* and *EDS1* were developed previously (Schultink et al., 2017). The constructed TRV2 vectors were transformed into *Agrobacterium tumefaciens* strain GV3101 and inoculated into 4-5 week-old *N. benthamiana* plants along with TRV1 at an OD600 of 0.2 for each. 2-3 weeks later, *Agrobacterium* transformed with different TNL/CNL and/or effectors were infiltrated into the VIGS plants for HR phenotype observation.

## Acknowledgments

T.Q. is supported by the Tang Distinguished Scholarship at the University of California, Berkeley. The work of A.S. was funded by US Department of Agriculture award UFDSP00011008. The Staskawicz lab is supported by the Two Blades Foundation.

## Supporting Information

**Supplemental Figure 1.**
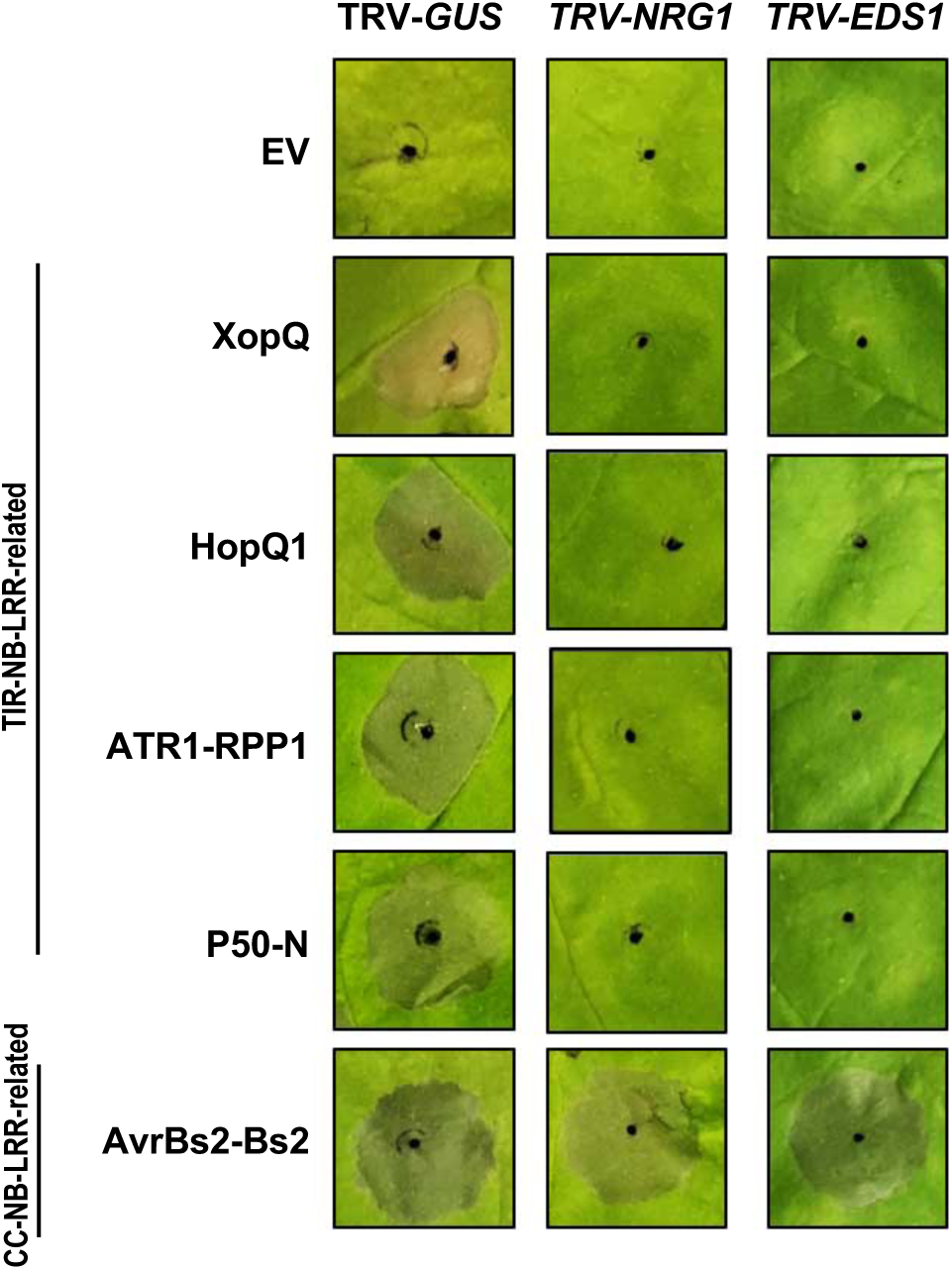
Virus-induced gene silencing (VIGS) of *NRG1* compromised several TNLs-mediated hypersensitive response in *N. benthamiana*. Tobacco rattle virus (TRV)-based VIGS was performed to down-regulate *GUS* (negative control), *NRG1* or *EDS1* in *N. benthamiana* leaves. Images of phenotypes of HR responses were taken 2 days after *Agrobacterium*-mediated transient expression of the empty vector control (EV), XopQ, HopQ1, ATR1-RPP1, N-P50 or AvrBs2-Bs2. The infiltrated leaves were wrapped up with aluminum foil for 2 days before images were taken.

**Supplemental Figure 2.**
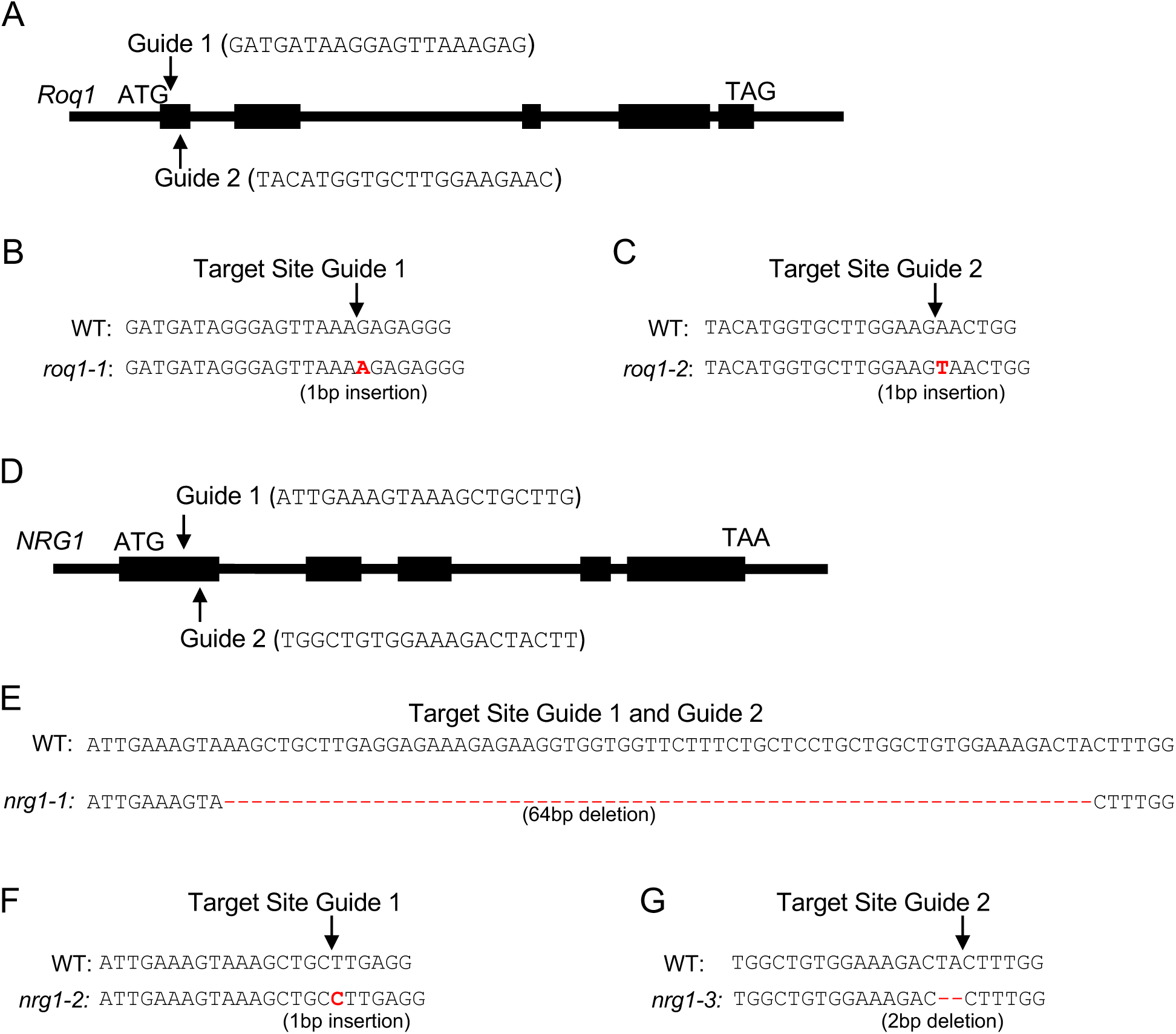
*Roq1* guides, *NRG1* guides and mutant alleles. (A) Two guides were designed to target the first exon of the *N. benthamiana Roq1* gene. (B and C) Two constructs containing Guide 1 or Guide 2 were transformed into *N. benthamiana* and regenerated plants were genotyped. Two frameshift alleles containing an A or T insertion at the target sites, named *roq1-1* (B) and *roq1-2* (C) respectively, were identified. The predicted cut site for each guide is indicated with an arrow and the inserted DNA bases causing a frameshift are in red and bold. (D) Two guides were designed to target the first exon of the *NRG1* gene in *N. benthamiana*. (E to G) The construct containing both Guide 1 and Guide 2 was transformed into *N. benthamiana* and the regenerated plants were genotyped. Three mutant alleles named *nrg1-1 (E), nrg1-2* (F), and *nrg1-3* (G) were identified. The predicted cut site for each guide is indicated with an arrow and the inserted DNA bases causing a frameshift are in red and bold.

**Supplemental Figure 3.**
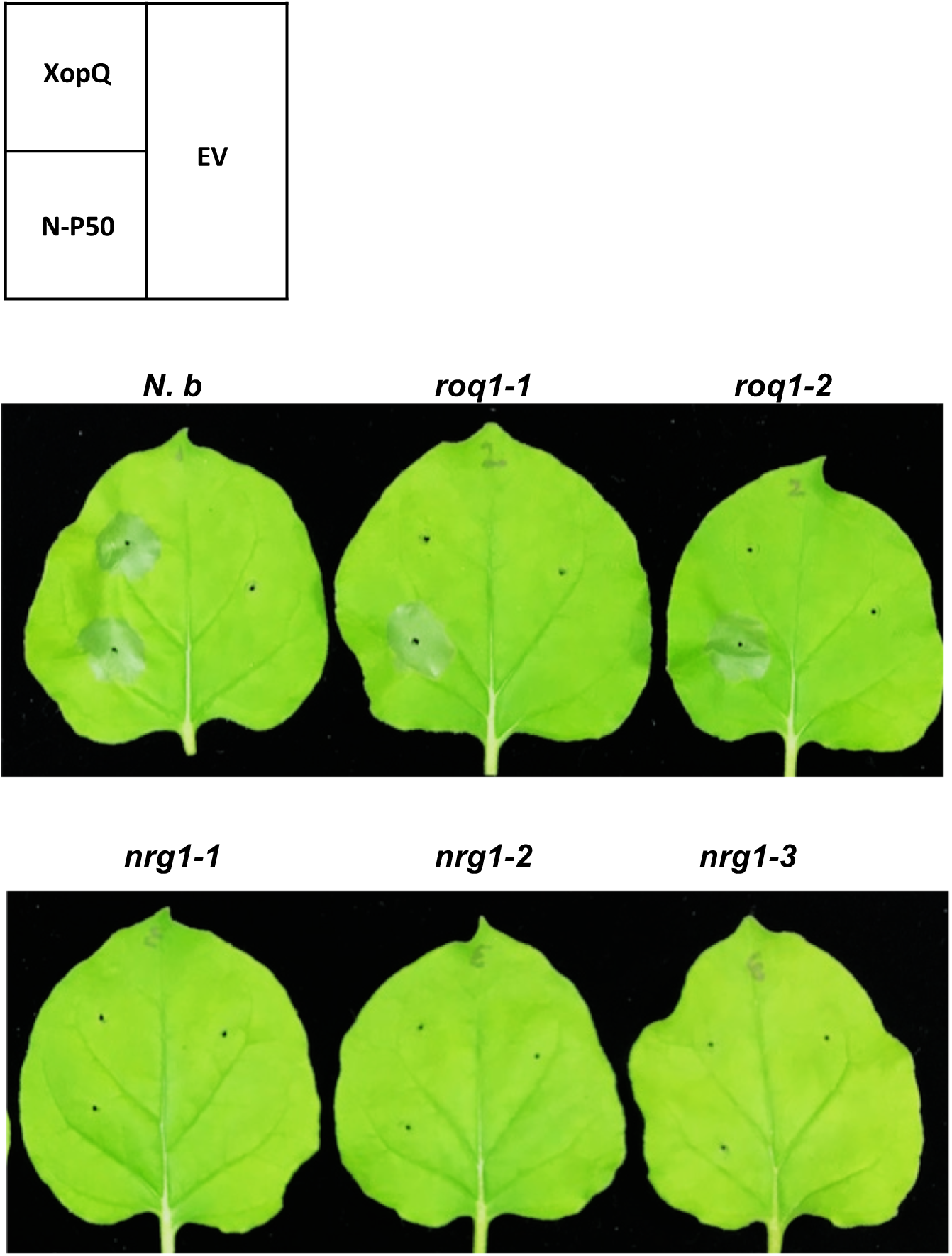
XopQ is unable to trigger hypersensitive response in the *nrg1* and *roq1* mutants. Hypersensitive response in leaves of *N. benthamiana* wild-type (*N. b*), *roq1-1, roq1-2, nrg1-1, nrg1-2* and *nrg1-3* with *Agrobacterium*-mediated transient expression of the empty vector control (EV), N-P50 or XopQ. The infiltrated leaves were wrapped up with aluminum foil and images were taken 2 days post-infiltration.

**Supplemental Figure 4.**
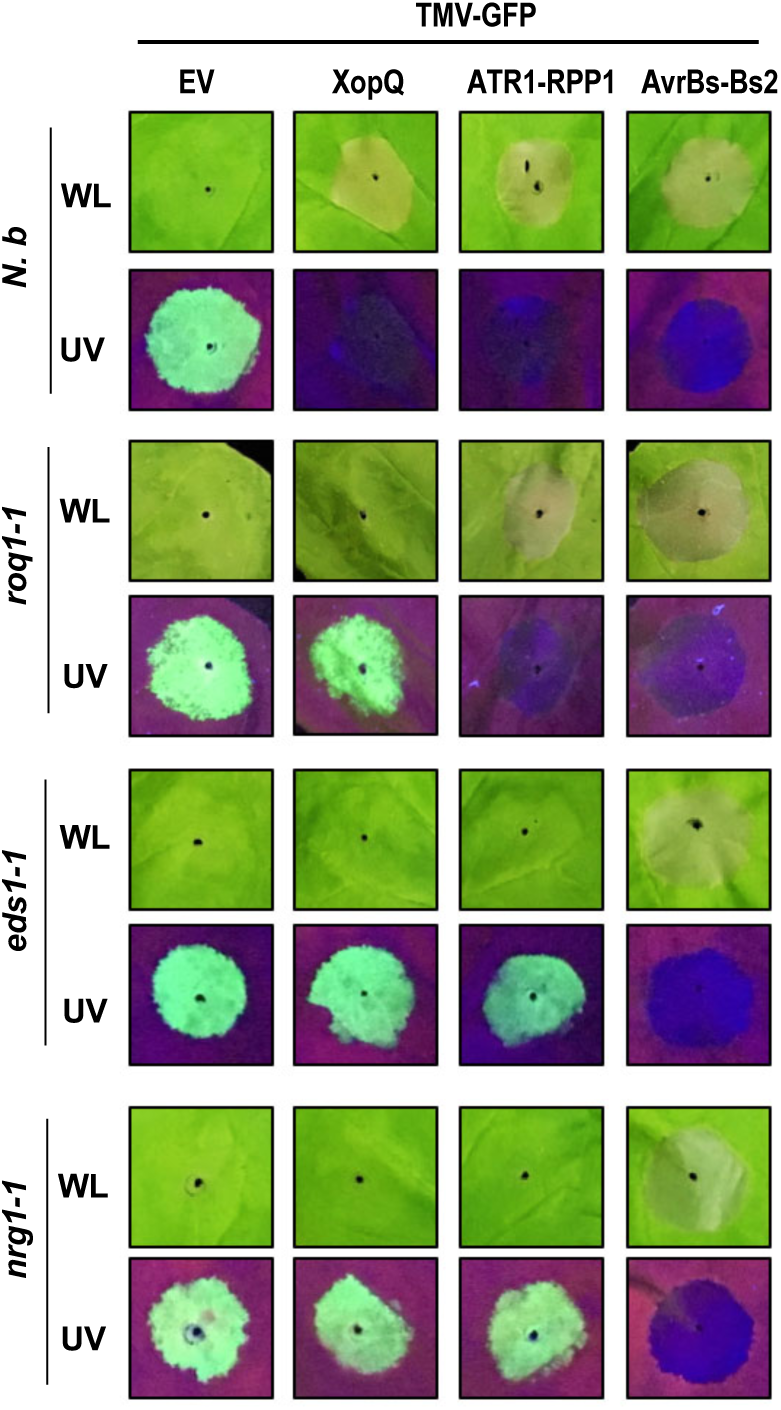
Roles of *NRG1* and *EDS1* in effector-activated defense against TMV. Phenotypes of HR reactions and TMV-GFP in leaves of *N. benthamiana* wild-type, *roq1-1, eds1-1*, or *nrg1-1* with *Agrobacterium*-mediated transient expression of TMV-GFP reporter plus the empty vector control (EV), XopQ, ATR1-RPP1, or AvrBs2-Bs2. The infiltrated leaves were wrapped up with aluminum foil and were imaged under white light (WL) or long-wave UV 3 days post-infiltration.

**Supplemental Table 1.**
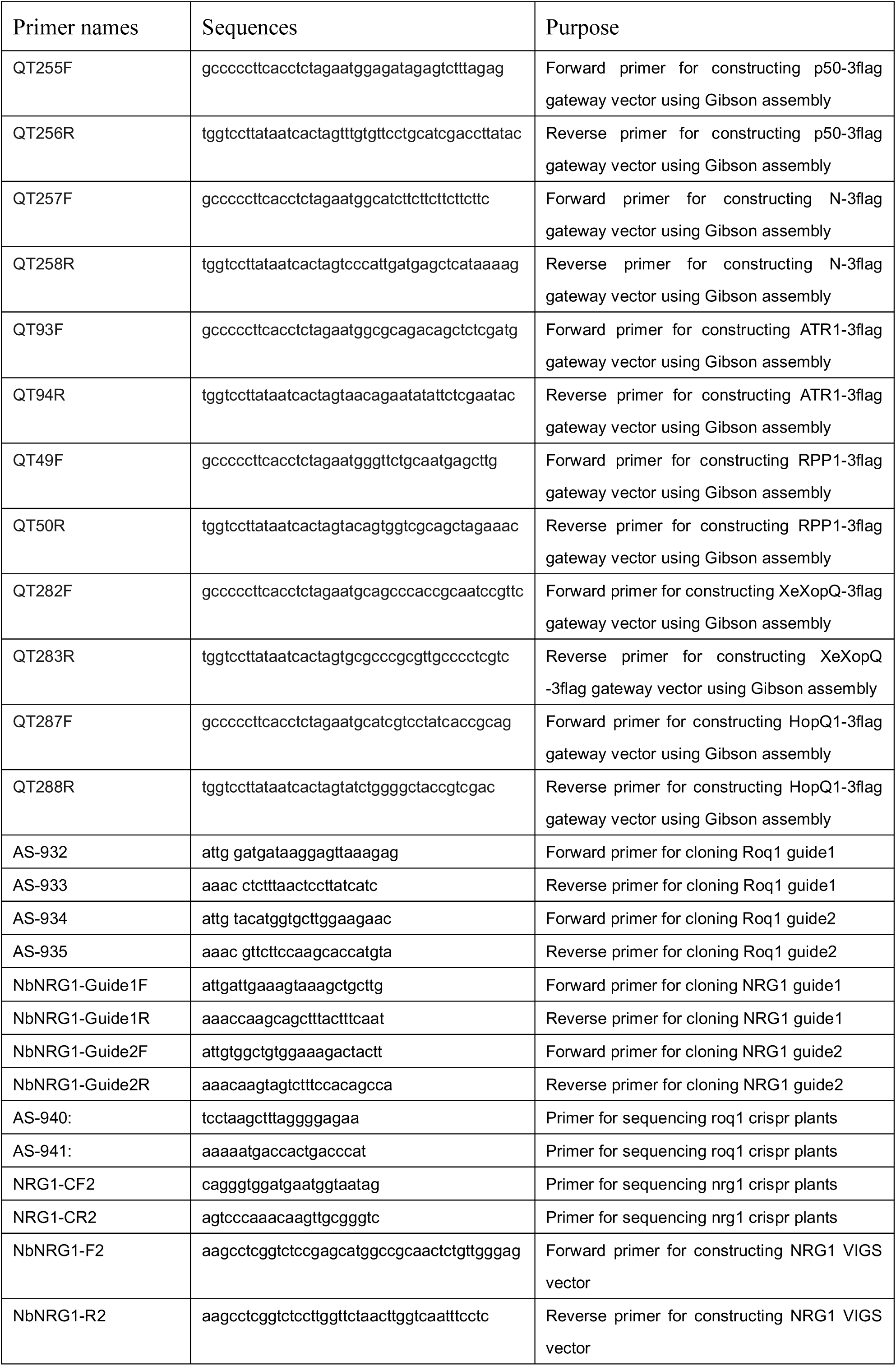
Primer sequences.

